# CIARA: a cluster-independent algorithm for the identification of markers of rare cell types from single-cell RNA seq data

**DOI:** 10.1101/2022.08.01.501965

**Authors:** Gabriele Lubatti, Marco Stock, Ane Iturbide, Mayra L. Ruiz Tejada Segura, Richard Tyser, Fabian J. Theis, Shankar Srinivas, Maria-Elena Torres-Padilla, Antonio Scialdone

**Author notes:** contributed equally (Marco Stock, Ane Iturbide).

## Abstract

A powerful feature of single-cell RNA-sequencing data analysis is the possibility to identify novel rare cell types. However, rare cell types are often missed by standard clustering approaches. We have developed CIARA (Cluster Independent Algorithm for the identification of markers of RAre cell types), a computational tool available in R and Python that outperforms existing methods for rare cell type detection. With CIARA, we found a small group of precursor cells among mouse embryonic stem cells and previously uncharacterized rare populations of cells in a human gastrula.

---

Characterizing rare cells has a fundamental importance in many biological contexts: for example, during development, to pin down at what stage a given cell type starts to emerge; when studying cancer, to look for rare cells that might develop drug resistance [1]; or for the characterization of stem cell lines, searching for cell transitions in different pluripotency states [2,3]. In addition to being rare, some types of cells can be challenging to identify because they have overlapping markers with other, more abundant cell types. This is the case, for instance, of primordial germ cells, which share markers with cells from the primitive streak [4,5].

Many algorithms specifically designed to detect rare cell types in scRNA-seq data have been devised (e.g., [6–10]). These methods generally work well in selecting rare cells with strong markers, but they are less efficient in identifying very small cell populations (<1%) with a limited number of unique markers. Moreover, some of these methods tend to overfit and identify a large number of small cell clusters without specific markers.

The algorithm we developed, CIARA, identifies potential marker genes of rare cell types by exploiting their property of being highly expressed in a small number of cells with similar transcriptomic signatures. To this aim, CIARA ranks genes based on their enrichment in local neighborhoods defined from a K-nearest neighbors (KNN) graph. The top-ranked genes have, thus, the property of being “highly localized” in the gene expression space. To identify potential groups of rare cell types, CIARA generates 2D representations of the data (eg, with UMAP [11]) showing how many and which of the top selected genes each cell expresses and shares with its neighbors (see Figure 1a, right panel; see also Methods and Supplementary Figure 1a-c for an example). Such a 2D map is also available in an interactive format, where the names of the genes are displayed for each selected cell (see examples in Supplementary Files 7-9). The identified top genes can be used with standard clustering algorithms to define the groups of rare cell types, either on the whole dataset or within specific clusters that were previously defined (see Figure 1a, right panel and Methods).

**Figure 1.**
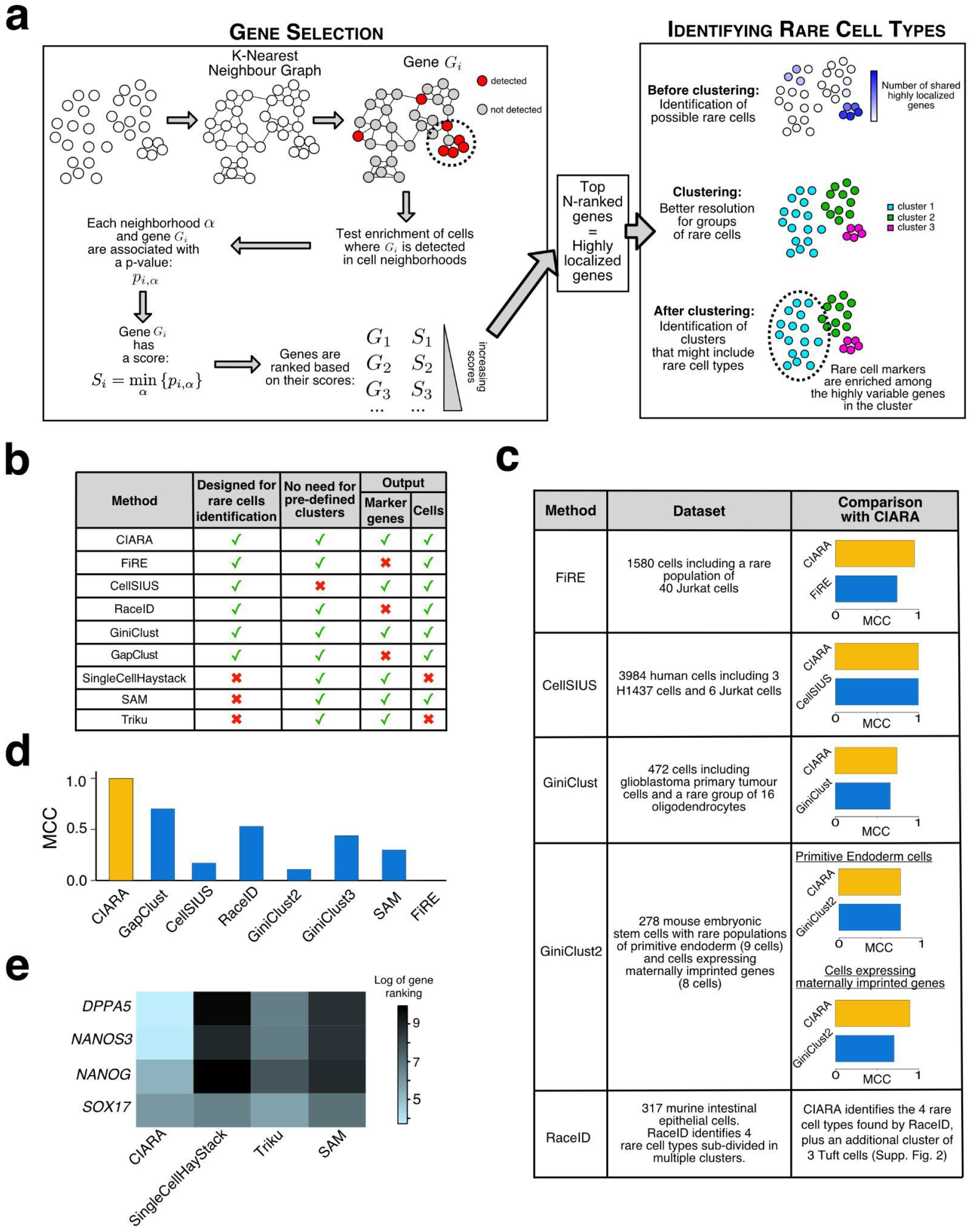
Schematic representation and benchmarking of CIARA. **a,** CIARA computes a score for each gene based on how cells expressing that gene are distributed on a K-nearest neighbor graph (left panel). Lower scores correspond to genes that are mostly expressed in neighboring cells, ie, are “highly localized”, and hence are more likely to be markers of rare cell types. The right panel summarizes how the top-ranked genes are used to visualize and identify groups of rare cells. **b,** Table listing the methods for rare cell-type identification we benchmarked CIARA against. We indicate particular features of each approach. **c,** Table summarizing the data sets and results of the benchmarking analysis. The last column shows the values of the Matthews Correlation Coefficient (MCC) computed between the group of rare cells identified by each method and the ground truth. **d,** MCC computed for the PGC group of cells present in the human gastrula data [4]. **e,** Heatmap showing the ranking (in natural log scale) of four PGC markers (in rows) obtained by four methods (in columns).

We tested the performance of CIARA against several existing methods to detect rare cell types: GiniClust [8,12], CellSIUS [6], FiRE [7], RaceID [9] and GapClust [10]. The features of the algorithms are summarized in Figure 1b. To make the comparison as fair as possible and minimize the effects of confounding factors due to, e.g., the use of sub-optimal parameter settings, we ran a first series of tests on the datasets included in the papers where the alternative algorithms were introduced. We evaluated performance by quantifying the agreement between the classification of rare cells obtained with each method and the ground truth classification using Matthew’s correlation coefficient (MCC; see Methods). The results of the benchmarking show that CIARA generally outperforms the other algorithms (see Figure 1c, Methods, and Supplementary Figure 2a-g).

We also ran all algorithms on the same recently published scRNA-seq dataset of 1,195 cells from a human gastrula [4], where a small population of 7 primordial germ cells (PGCs) is included. In addition to being rare, PGCs have markers in common with other cell types, like *SOX17* and *ETV4*, which complicates the identification of PGCs with unsupervised methods. CIARA detects a cluster including all the 7 PGCs, achieving an MCC = 1 (see Methods). Conversely, all other algorithms achieved a lower MCC value (Figure 1d, Methods and Supplementary figure 2h-i). In addition to the algorithms mentioned above, we ran three more algorithms on the human gastrula dataset: singleCellHayStack [13], SAM [14], and Triku [15]. Although not specifically designed for detecting rare cell types, these algorithms find genes that have a non-random distribution of expression values across cells. While these approaches offer a valid alternative to standard differential expression analysis methods, they tend to miss rare cell markers, as is seen with PGC markers (see Figure 1e and Methods).

Overall, these analyses show that CIARA has better performance than alternative algorithms to detect rare cells in several published datasets, also in the most challenging situations when the rare cells share marker genes with more abundant cell types.

To show how CIARA can produce new biological insights, we applied it to two datasets where cell differentiation occurs.

*In vitro* differentiation systems are particularly suitable to study how, for example, signaling pathways can regulate fate decisions. Recently, we showed that a 48h-long treatment with low doses of retinoic acid (RA) induces the reprogramming of mouse embryonic stem cells (mESCs) into 2-cell-like cells (2CLC) [16], a cell type that resembles totipotent cells [3]. However, it is not known how long the RA treatment must be to produce any effects on cell fate decisions. Thus, we generated a new scRNA-seq dataset from mESC following a 24h RA treatment (Figure 2a and Supplementary Figure 3a-d) and we analyzed the data with CIARA looking for changes in cell type composition. In addition to a cluster of pluripotent cells (744 cells, ~97%) and 2CLC (18 cells, ~2%), CIARA also detected a small group of 4 cells (<1%) marked by a distinct set of genes, including differentiation markers like *Gata4* and *Gata6* (Figure 2b-c, Supplementary File 1 and 8). A comparison with previously published datasets confirmed that this small cluster includes 4 precursor cells that are compatible with those found at 0h and 48h of treatment (see Methods). These results indicate that during the first 48h of RA treatment, the same cell types are present, while there is a shift in their relative abundance only after more than 24h of treatment (Supplementary Figure 3e).

**Figure 2.**
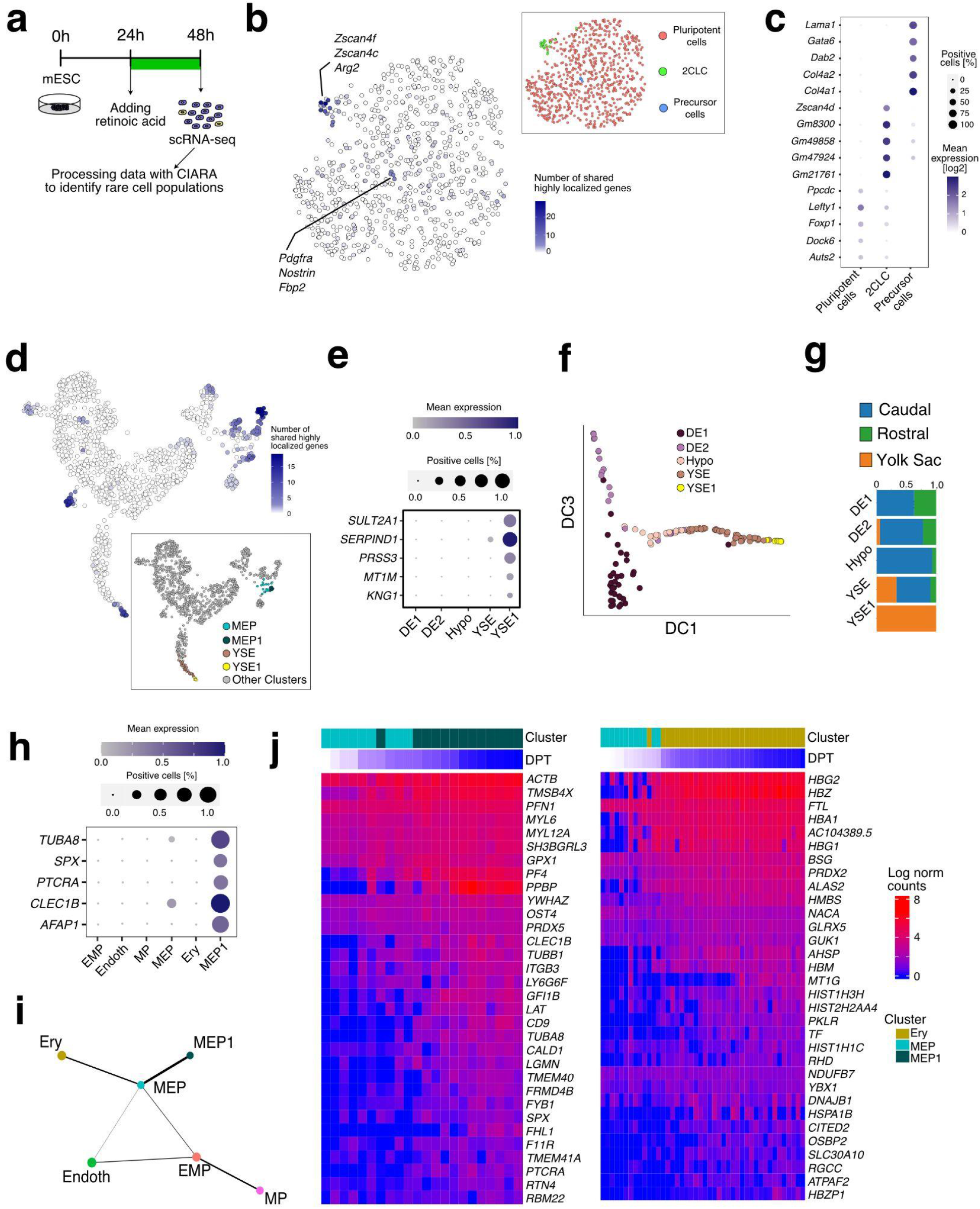
CIARA identifies rare populations of cells in mESC and a human gastrula dataset. **a,** We treated mESC with retinoic acid for 24h before collecting and processing them for scRNA-seq. **b,** UMAP representation of the mESC dataset indicating the number of highly localized genes expressed by each cell and shared with their neighbours. Inset: cells are colored by cluster. **c,** Top marker genes of the clusters found in the mESC data. **d,** UMAP representation of the human gastrula dataset [4] showing the number of shared highly localized genes in each cell. Inset: The sub-clusters highlighted, YSE and MEP, are those in which CIARA finds new rare cell populations (YSE1 and MEP1). **e,** Top marker genes of the YSE1 rare cell population. Mean expression levels are normalized by the maximum within each cluster. **f,** Diffusion components 1 and 3 (DC1, DC3) of the endodermal cells. **g,** Stacked barplot showing the distribution of the anatomical origin of cells in each cluster. **h,** Top marker genes of the MEP1 rare cell population. Mean expression levels are normalized by the maximum within each cluster. **i,** Graph representation of the connectivity between the clusters of blood cells estimated with PAGA [21]. **j,** Top differentially expressed genes along the differentiation trajectories joining MEP and MEP1 (left panel) or MEP and Ery (right panel). YSE, Yolk Sac Endoderm; MEP, Megakaryocyte-Erythroid Progenitors; Ery, erythroblasts; Endoth, endothelium; EMP, erythro-myeloid progenitors; MP, myeloid progenitors; DE, Definitive Endoderm; Hypo, Hypoblast.

Finally, we applied CIARA to a previously published scRNA-seq dataset from a human gastrula [4]. Using CIARA, we performed an unsupervised analysis of this dataset looking for rare cell types. In addition to the PGCs described above (Figure 1d-e), we found two small populations in the Yolk Sac Endoderm (YSE) and the Megakaryocyte-Erythroid Progenitors (MEP) clusters (Figure 2d and Supplementary File 9). The small YSE subcluster of 11 cells that we named YSE1 expresses very specific markers (e.g., *SERPIND1, SERPINC1;* Figure 2e, Supplementary Figure 4b and Supplementary File 4). These genes are known to be expressed in the adult kidney and liver [17], which is consistent with the functions that the yolk sac plays during early development [18]. Interestingly, CIARA identifies a small cluster of 21 endodermal cells with the same transcriptional profile in mouse embryos at the E7.75-E8.25 stage ([19]; see Methods, Supplementary Figure 4d and Supplementary File 6). This observation indicates that YSE1 is a relatively rare endodermal sub-population present in human and mouse embryos. A diffusion map and pseudo-time analysis of the human Endoderm cluster reveal that YSE1 is more transcriptionally distinct from the embryonic endoderm populations (represented by the Definitive Endoderm clusters) than the rest of the YSE cluster (Figure 2f and Supplementary Figure 4c). Furthermore, all cells included in YSE1 come from the yolk sac region, while the rest of the YSE cluster also includes cells from the embryonic disk (Figure 2g). This suggests that YSE1 might represent a cell population in the yolk sac endoderm located further away from the embryonic disk, and possibly closer to the forming blood islands where primitive erythropoiesis occurs [4]. In support of this hypothesis, one of the markers of YSE1 is transferrin (*TF*), a protein iron carrier fundamental for erythropoiesis [20] whose receptors *TFRC* and *TFR2* are expressed by erythroblasts (Supplementary Figure 4e).

The second population of rare cells detected by CIARA is in the Megakaryocyte-Erythroid Progenitor (MEP) cluster, which we named MEP1 (Figure 2d). This cluster is made of 13 cells with a distinct transcriptional signature characterized by high levels of markers like *PPBP, ITGA2B*, and *GP1BB* (Figure 2h, Supplementary Figure 4f, and Supplementary File 5). Based on these and other markers, we could identify these cells as megakaryocytes. This conclusion is also supported by the analysis of the differentiation trajectories within the blood clusters (Figure 2i-j and Methods). Specifically, we found a branching event where the MEP cluster splits into the MEP1 cluster (where megakaryocytes markers are upregulated) and erythroblasts (Figure 2i), allowing us to identify genes marking the differentiation between these two cell types (Figure 2j). An analogous rare population of megakaryocytes with the same transcriptional signature was also identified in mice at a later developmental stage (see Methods), which is consistent with human hematopoiesis starting earlier than in mice, as suggested by other analyses [4].

These applications exemplify two general tasks for which CIARA can be employed: first, the detection of small changes in cell type composition over a time course experiment; and the characterization of a system where new cell types are just emerging, to pinpoint the first transcriptional steps that accompany cellular fate decisions.

While CIARA is a powerful method to identify and characterize rare cell types, it relies on very few parameters (see Methods) given that it uses an exact probability distribution to compute the gene scores and does not need pre-defined clusters (similarly to recently published methods to compare cell type abundance across conditions [22]). Moreover, its main requirement is simply the definition of a KNN graph. Hence, its applicability can be easily extended to other single-cell data, such as ATAC-seq. This would allow the identification of rare cells across multiple modalities, which could help validate the presence of new rare cell populations and lead to a more in-depth characterization of cell types as they differentiate.

## Methods

### CIARA algorithm

#### Gene selection

CIARA starts from a normalized gene count matrix and K-nearest neighbor (KNN) graph, which can be built with standard approaches available in the seurat [23] or scanpy [24] libraries. Given its goal to find potential markers of rare cells, CIARA performs a filtering step to select only genes that are expressed above a threshold value in a relatively small number of cells. The thresholds can be set manually or the default values will apply (threshold expression value: threshold=1; minimum number of cells: n_cells_low=3; maximum number of cells: n_cells_high=20).

For the genes that pass this filtering step, CIARA will carry out a one-sided Fisher’s exact test to check whether there is a statistically significant enrichment of cells expressing the gene in each neighborhood (formed by a cell and its KNN). This is done with the function “fisher.test” in R, with the option “alternative=greater”. By default, the result of the test is considered statistically significant if the p-value is smaller than 0.001.

All the genes that show statistically significant enrichment in at least one neighborhood have expression patterns that are highly localized and are considered potential markers of rare cell types. These highly localized genes are assigned a score equal to the minimum p-value obtained across all neighborhoods (Figure 1a). Such score is used to rank the genes, with smaller scores being associated with genes that are more strongly enriched in at least one neighborhood.

If a gene is not enriched in any neighborhood, it gets assigned a score equal to 1.

#### Identifying rare cells

The highly localized genes (having a score < 1, see above) are used by CIARA to identify groups of rare cell types following two main strategies.

The first is clustering-independent, and consists of counting for every single cell the number of highly localized genes expressed in that cell and in its KNN: the larger this number, the more likely it is that the cell is part of a group of rare cells. The results of this analysis are reported in a 2D representation of the data, like a UMAP plot (see Figure 2b,d, and Supplementary Figure 1a), where each cell is colored based on the number of highly localized genes expressed and shared across the KNN. Such 2D plot is also available in an interactive html format, where by hovering the mouse cursor over any cell, the names of the top highly localized genes expressed and shared across the KNN are shown (see Supplementary File 7,8 and 9).

The second strategy for rare cell type identification is based on utilizing standard clustering algorithms with the highly localized genes selected by CIARA. In the R version of CIARA, clustering is done with the Louvain algorithm on the first 30 principal components as a default value (defined from the top 2000 highly variable genes) with the functions FindNeighbors and FindClusters from the R library seurat version 4.0.5.

The clustering can involve the entire dataset or only part of it. In particular, given an existing partition of the data, CIARA can verify which clusters are more likely to include groups of rare cell types by testing the enrichment of highly localized genes among the top 100 (default value) highly variable genes within each cluster (Fisher’s test, p-value < 0.001 and odds ratio greater than 1). Those clusters that show a significant enrichment are then sub-clustered with the same algorithm as specified above.

#### Marker genes identification

The markers for the clusters identified by CIARA are detected with the FindMarkers function (with parameter only.pos = T) from seurat (version 4.0.5). Only markers with adjusted p-value (based on the Bonferroni correction) below or equal to 0.05 are considered for downstream analysis. Finally, for each cluster, only unique markers (ie, that are not included among the markers of other clusters) are kept.

Unless otherwise specified, in the balloon plots showing marker genes expression the size of the dots is determined by the fraction of cells with log norm counts above 1 (function NormalizeData from R library seurat).

### Analysis of previously published datasets for benchmarking analysis

Below, we briefly describe the datasets we used for the benchmarking analysis shown in Figure 1c. To evaluate the performance of each algorithm, we quantified the agreement between the classification of rare cells obtained with each method and the ground truth classification using the Matthew correlation coefficient (MCC). MCC is a metric that quantifies the overall agreement between two binary classifications, taking into account both true and false positives and negatives. MCC values range from (−1) to 1, where 1 indicates a perfect agreement between clustering and the ground truth, 0 means the clustering is as good as a random guess, and (−1) indicates no overlap between the clustering and the ground truth. MCC is computed with the function mcc from the R library mltools version 0.3.5 (https://CRAN.R-project.org/package=mltools). The MCC values shown in Figure 1c-d for each algorithm represent the maximum values obtained across all clusters.

In all the datasets analyzed with CIARA, the normalized count matrix was obtained with the function NormalizeData (with parameter normalization.method = LogNormalize) and the K-nearest neighbor (KNN) graph was built with the function FindNeighbors (on the first 30 principal components built from the top 2000 highly variable genes). Both functions are from seurat version 4.0.5.

#### Dataset: 293T and Jurkat cells (Figure 1c)

This dataset of 1580 cells comprises 293T and Jurkat cells in a known proportion, with the Jurkat cells being the rare population (40 cells, ~2.5% of total cells). This dataset was analyzed in the FiRE paper [7].

Here, CIARA identified 2077 highly localized genes. By clustering the data with these genes, we found 2 clusters (resolution 0.1, k.param equal to 5 and number of principal components equal to 30), one of which corresponded to Jurkat cells, based on the markers expressed.

CIARA outperforms FiRE (MCC values are 0.95 and 0.74 respectively; see Figure 1c), due to the fewer false positives (4 cells) compared to those detected by FiRE (32 cells).

#### Dataset: mixture of eight human cell lines (Figure 1c)

This dataset includes 3984 cells, and was analyzed in the CellSIUS paper [6]. Two rare populations of H1437 and Jurkat cells (3 and 6 cells respectively) are present and marked in the dataset.

Here, CIARA identified 3704 highly localized genes. By clustering the data with these genes, we identified 9 clusters (resolution 0.1, k.param equal to 3, and number of principal components equal to 30). Two of these clusters could be identified as H1437 and Jurkat cells based on their markers. Hence, CIARA could identify both of these rare cell types, achieving the same performance as CellSIUS (MCC equal to 1 for both methods; Figure 1c).

#### Dataset: glioblastoma (GBM) primary tumors (Figure 1c)

This dataset includes 472 cells, and was analyzed in the GiniClust paper [25]. It includes a small group of 16 oligodendrocytes, which are defined as the cells co-expressing the four marker genes *CLDN11, MBP, PLP1*, and *KLK6* [25]. CIARA identified 68 highly localized genes. By clustering the data with these genes, we identified 13 clusters (resolution 0.1, k.param equal to 3 and number of principal components equal to 30), one of which corresponded to oligodendrocytes.

#### Dataset: differentiating mouse embryonic stem cell (mESC) at day 4 post LIF-withdrawal (Figure 1c)

This dataset includes 278 mESC that are differentiating after LIF removal, and was analyzed in GiniClust2 paper [12]. On day 4 after LIF removal, two small clusters of cells (9 and 8 cells) are detected by GiniClust2, which, based on their markers, were identified as cells differentiating towards primitive endoderm (PrE cells; markers: *Col4a1, Col4a2, Lama1, Lama2*, and *Ctsl*) and cells expressing maternally imprinted genes (*Rhox6, Rhox9*, and *Sct*).

CIARA identified 287 highly localized genes. By clustering the data with these genes, we identified 3 clusters (resolution 0.3, k.param equal to 5 and number of principal components equal to 30). Two of these clusters expressed the same markers as the rare cells identified by GiniClust2 (Supplementary figure 2a-c). While in this dataset we lack a “ground truth” for the rare cells, we define a set of “bona fide” clusters based on the co-expression of the marker genes mentioned above, and we compute the MCC values of GiniClust2 and CIARA using these clusters as reference. The two methods had the same MCC score for the cluster of differentiating cells, but CIARA achieved a higher MCC value on the set of cells expressing maternally imprinted genes (Figure 1c).

#### Dataset: murine intestinal epithelial cells (Figure 1c)

This dataset includes 317 cells and was analyzed in the RaceID [26] vignette (https://cran.r-project.org/web/packages/RaceID/vignettes/RaceID.html). Here, 4 rare cell types (enterocytes, goblet cells, Paneth cells, and enteroendocrine cells) are found after manually merging multiple clusters expressing similar marker genes [26]. CIARA identified 1514 highly localized genes. By clustering the data with these genes, we found 8 clusters (resolution 0.2, k.param equal to 3, and number of principal components equal to 20), 4 of which correspond to the rare cell types that RaceID finds. Additionally, one of the clusters found by CIARA (cluster number 7) expresses markers of Tuft cells (Supplementary Figure 2f).

The markers for the dataset (using the clusters defined with CIARA, see Supplementary Figure 2 d-f) are identified as specified above.

To investigate the relationship between the 6 smallest clusters (<= 25 cells) detected by CIARA(2, 3, 4, 5, 6, and 7) and the original cluster partition obtained with RaceID, a plot was generated with the function clustree from the R package clustree version 0.4.4 ([27]; Supplementary Figure 2g).

Among these clusters identified by CIARA, cluster 2 corresponds to Goblet cells (marked by *Clca3*), cluster 3 to enterocytes (marked by *Apoa1*), cluster 4 to Paneth cells (marked by *Defa24*), cluster 5 to enteroendocrine cells (marked by *Chgb*), and cluster 7 to Tuft cells [9]. The markers used to label the clusters from 2 to 5 were described in Supplemental Information Figure S2 B from [26].

The clustree plot in Supplementary Figure 2g shows that each of the above rare cell types identified by CIARA are split between several clusters with RaceID.

Cluster 6 (4 cells) shows a very clear transcriptional profile and corresponds to a cell type not previously described (Supplementary figure 2e).

### Identification of Primordial Germ Cells from a human gastrula dataset

We analyzed a previously published human gastrula dataset from [4] using CIARA and the other seven algorithms we tested in Figure 1. Among the 1195 cells of this dataset, there is a small population of seven Primordial Germ Cells (PGCs), which were identified in [4] in a supervised way (ie, by using the co-expression of known PGC markers like *NANOS3, NANOG*, and *DPPA5*). We describe below how we ran the algorithms and tested their ability to find PGCs.

CIARA found 2917 highly localized genes in the whole dataset. By clustering the data with these genes, the seven PGCs are always identified as a single cluster over a wide range of resolutions (Supplementary Figure 4b).

GiniClust2 and GiniClust3 pipelines were used following the documentation available from https://github.com/dtsoucas/GiniClust2 and https://github.com/rdong08/GiniClust3 with default values for all parameters. Note that the gene selection based on the Gini index tends to miss PGC markers due to their low average expression values (Supplementary Figure 2h-i).

For CellSIUS, we used the R package available from https://github.com/Novartis/CellSIUS/. We decreased the value of the “min_n_cells” parameter from its default value (10) to 5 (given that there are only seven PGCs in the data), while default values were used for the other parameters. The FiRE R package is available from https://github.com/princethewinner/FiRE. Using the default threshold on the FiRE score (i.e., 1.5*interquartile range + third quantile), no rare cells were identified. Hence, we chose a less stringent threshold of 0.5*interquartile range + third quantile. Since FiRE does not provide clusters of cells as output, for the MCC computation we considered the rare cells identified by FiRE in the Primitive Streak cluster as PGCs.

The analysis with RaceID 3 was performed with standard parameters using the R package https://github.com/dgrun/RaceID3_StemID2_package.

For the analysis with GapClust, we used the implementation available from the GitHub repository https://github.com/fabotao/GapClust with default parameters.

The SingleCellHaystack algorithm is implemented in the R package available from https://github.com/alexisvdb/singleCellHaystack. Default values were used for all parameters, and the algorithm was run on the first thirty principal components.

Analysis with SAM was performed with default values of all parameters from the Python package https://github.com/atarashansky/self-assembling-manifold/tree/master.

The Triku algorithm is implemented from the Python package available from the website https://triku.readthedocs.io/en/latest/. This website also includes a tutorial that we followed to perform our analysis. For gene filtering, we ran the function pp.filter_genes from scanpy (verions 1.8.0) with min_cells = 3 instead of the default value equal to 10 (given that the number of PGC is less than 10).

SingleCellHaystack, SAM, and Triku return a ranked list of “most informative” genes having a non-random distribution of expression values across cells. We verified whether the top 1000 genes selected by these three algorithms were enriched with PGC markers by running a Fisher’s test (R function fisher.test with alternative = “two.sided”) using as background all the genes with normalized expression above 0.5 in more than 6 cells. None of the tested methods showed a statistically significant enrichment of PGC markers, apart from CIARA (p-value = 8*10^-4^).The data normalization is done with the function NormalizeData from Seurat with parameter normalization.method = “LogNormalize”.

The PGC markers were detected with the FindMarkers function (with parameter only.pos = T) from Seurat using a threshold for the Bonferroni-adjusted p-value of 0.05 and excluding all genes that were also markers of other non-PGC clusters.

### Mouse embryonic stem cells experiment

#### Cell culture

Cells were grown in a medium containing DMEM-GlutaMAX-I, 15% FBS, 0.1mM 2-beta-mercaptoethanol, non-essential amino acids, penicillin and streptomycin and 2× LIF over gelatin-coated plates. The medium was supplemented with 2i (3 μM CHIR99021 and 1 μM PD0324901, Miltenyi Biotec) for maintenance and expansion. The 2i was removed 24h before the addition of retinoic acid (RA) as described in Iturbide et al 2021 [16].

#### Single-cell RNA-seq

Cells were collected after RA treatment and sorted for live single cells by FACS. Cells were then counted and tested for viability with an automated cell counter. Five thousand cells of the sample were then input into the 10X protocol. Gel bead-in-emulsion (GEM) generation, reverse transcription, cDNA amplification, and library construction steps were performed according to the manufacturer’s instructions (Chromium Single Cell 3’ v3, 10X Genomics). Samples were run on an Illumina NovaSeq 6000 platform.

#### Gene counting

Unique molecular identifier (UMI) counts were obtained using the kallisto (version 0.46.0) bustools (version 0.39.3) pipeline [28]. First, the mouse transcriptome and genome (release 98) fasta and gtf files were downloaded from the Ensembl website, and 10X barcodes list version 3 was downloaded from the bustools website. We built an index file with the ‘kallisto index’ function with default parameters. Then, pseudoalignment was done using the ‘kallisto bus’ function with default parameters and the barcodes for 10X version 3. The BUS files were corrected for barcode errors with ‘bustools correct’ (default parameters), and a gene count matrix was obtained with ‘bustools count’ (default parameters).

#### Quality control and normalization

To remove barcodes corresponding to empty droplets, we used the ‘emptyDrops’ function from the R library ‘DropletUtils’ version 1.6.1 [29]. For this, a lower threshold of 1,000 UMI counts per barcode was considered. Afterward, quality control was performed using the scanpy library. Cells having more than 10% counts mapped to mitochondrial genes or fewer than 1,000 detected genes were removed. After quality control, 766 cells were kept for downstream analysis (Supplementary Figure 3a-d).

#### Analysis with CIARA

CIARA identifies 2475 highly localized genes in this dataset. We ran cluster analysis on these genes with the FindNeighbors (on the 30 top principal components and with k.param equal to 3) and FindClusters functions (with resolution 0.1), which gave 3 clusters.

The marker genes of these clusters (see Supplementary File 1-3) were detected with the FindMarkers function (with parameter only.pos = T) from seurat. Only markers with an adjusted p-value based on Bonferroni correction below or equal to 0.05 (for 2CLC and Precursor cells) or with a p-value below 0.05 (for Pluripotent cells) are considered for downstream analysis. Moreover, for each cluster, only unique markers (eg, those not included in the marker list of other clusters) were kept.

Based on the lists of marker genes, the three clusters could be identified as Pluripotent Cells, 2-cell-like cells (2CLC), and Precursor Cells (Figure 2b-c).

#### Comparison with previously published mESC data

We compared the clusters found in our mESC dataset with those in the previously published mESC datasets after a 0h and 48h RA treatment [16]. The dataset at 0h was re-analyzed with CIARA, which identified 3302 highly localized genes. Using these genes, we performed clustering with the functions FindNeighbors (on the top 30 principal components with k.param = 5) and FindClusters with resolution 0.1, which gave three clusters. Based on their markers (found with the procedure described above), these clusters could be identified as Pluripotent Cells (1245 cells), 2CLC (36 cells), and Precursor Cells (4 cells; Supplementary Figure 1). These same clusters were identified in the dataset at 48h in [16].

We assessed the statistical significance of the intersection between the markers of the three clusters found at 0h, 24h, and 48h by using a Fisher’s test (with the fisher.test function from the R package stats, with ‘alternative=two.sided’).

The intersection between the markers of the Precursor Cells clusters at 24h versus 48h was significant with a p-value = 7*10^-48^; and at 24h versus 0h was significant with a p-value = 10^-31^. Similarly, the markers of the 2CLC cluster has a significant overlap at 24h versus 48h (p-value=9*10^-102^) and at 24h versus 0h (p-value = 6*10^-83^).

Finally, also the intersections between the markers of pluripotent cells at 24h versus 0h (p-value = 0.0001) and at 24h vs 48h (p-value = 2 * 10^-91^) are statistically significant.

### Identifying rare cell types in the human gastrula dataset

First, we tested the enrichment of the 2917 highly localized genes found by CIARA among the top 100 highly variable genes (HVGs) within each of the clusters provided in [4] (as described above in the section “CIARA algorithm - Identifying rare cells”).

We found a statistically significant overlap in the Endoderm (Endo; p-value = 4*10^-5^) and the Haemato-Endothelial Progenitors cluster (HEP; p-value=4*10^-5^). Then, we sub-clustered the Endo and the HEP clusters using their HVGs (for the Endo cluster: resolution = 0.2, k.param = 5, top 30 principal components; for the HEP cluster: resolution = 0.6, k.param= 5, top 30 principal components). The two smallest clusters found in the Endo and HEP clusters are denoted as YSE1 and MEP1 respectively, and they were not described in [4].

#### Markers analysis

The markers for the human gastrula are detected with the FindMarkers function (with parameter only.pos = T) from seurat, with the same criteria described above. The analysis was run separately using all sub-clusters reported in [4] for the Endo cluster (including the new rare cluster found by CIARA, YSE1) and the HEP cluster (including MEP1 found by CIARA).

#### Trajectory and PAGA analysis

For the cells in the Endo sub-clusters (i.e., DE1, DE2, YSE, Hypoblast, and YSE1), a diffusion map was computed from the normalized count matrix with the top 2000 highly variable genes (using the NormalizeData and FindVariableFeatures functions from Seurat) with the function DiffusionMap from the R package destiny version 3.2.0 [30]. The diffusion pseudo time is computed using the function DPT from the same package.

For the cells in the HEP sub-clusters (EMP, HE, MP, MEP, MEP1) and the Erythroblast cluster, the trajectory analysis was performed using the function slingshot (with start.clus=“MEP” and reducedDim equal to the diffusion map provided in the original human gastrula paper) from the R library slingshot version 1.6.1 [31].

To identify differentially expressed genes along the differentiation trajectories joining MEP and MEP1 or MEP and Ery, the functions fitGAM and startVsEndTest (with parameter lineage equal to TRUE) from the R package tradeSeq version 1.2.1 [32] were used.

To estimate the connectivity between clusters, we performed an analysis with PAGA [21] (functions tl.paga and pl.paga from scanpy).

#### Comparison with published mouse datasets

We analyzed with CIARA a previously published dataset from mouse embryos at E7.75-E8.25 stage [19]. This dataset of 665 cells included two small endodermal clusters. CIARA found 1700 highly localized genes (with *n_cells_high* = 30); using these genes for clustering (resolution = 0.2, k.param = 5 and number of principal components equal to 30), we could identify three clusters (Supplementary figure 4d), one of which is a small sub-cluster of 21 cells in the endodermal cluster labeled as “En2”. The markers of this sub-cluster (found with the procedure described above; Supplementary File 6) have a statistically significant overlap with the markers of the YSE1 cluster in the human gastrula (p-value = 0.0009, two-sided Fisher’s test).

In [5] a cluster of megakaryocytes was identified in mouse embryos. We tested the statistical significance of the overlap between the markers of these cells in mouse (from “source data figure 3f” in [5]) and the markers of MEP1 cluster from the human gastrula using a two-sided Fisher’s test, and obtained a p-value = 9*10^-7^).

The genes in the two mouse datasets ([19] and [5]) were converted into the corresponding human orthologous name if there is a 1:1 correspondence between the mouse and the human gene name, using g:Profiler [33].

## Data availability

Raw data for the mouse embryonic stem cells scRNA-seq data are available through ArrayExpress, under accession numbers EMTAB-11610.

## Code availability

The code used to generate the figures in this paper is available at https://github.com/ScialdoneLab/CIARA. In this repository, there are also additional examples of applications of CIARA.

CIARA is available both in R (https://CRAN.R-project.org/package=CIARA) and Python (https://github.com/ScialdoneLab/CIARA_python). Both packages can be easily integrated with standard analysis pipelines based on, e.g., Seurat [23] and Scanpy [24].

## List of Supplementary Files

Supplementary file 1: markers of precursor cells found in mESC treated with RA for 24h

Supplementary file 2: markers of 2CLC found in mESC treated with RA for 24h

Supplementary file 3: markers of pluripotent cells found in mESC treated with RA for 24h

Supplementary file 4: markers of YSE1 cluster in the human gastrula

Supplementary file 5: markers of MEP1 cluster in the human gastrula

Supplementary file 6: markers of sub-cluster of mouse endodermal cells equivalent to YSE1

Supplementary file 7: interactive plot for mouse ESCs treated with RA at 0h

Supplementary file 8: interactive plot for mouse ESCs treated with RA at 24h

Supplementary file 9: interactive plot for human gastrula dataset

## Acknowledgments

Work in the Scialdone lab is funded by the Helmholtz association. Work in the Torres-Padilla laboratory is funded by the Helmholtz Association, HMGU Small Molecule projects (Developmental projects) and the German Research Council (CRC 1064). A.I. was a recipient of a long-term EMBO fellowship (ALTF 383-2016). G.L. was funded by the BMBF project MechML (grant n. 01IS18053A). We thank members of the Scialdone lab for discussions and feedback on the manuscript.

