## Supplementary figures for "CIARA: a cluster-independent algorithm for the identification of markers of rare cell types from single-cell RNA seq data"

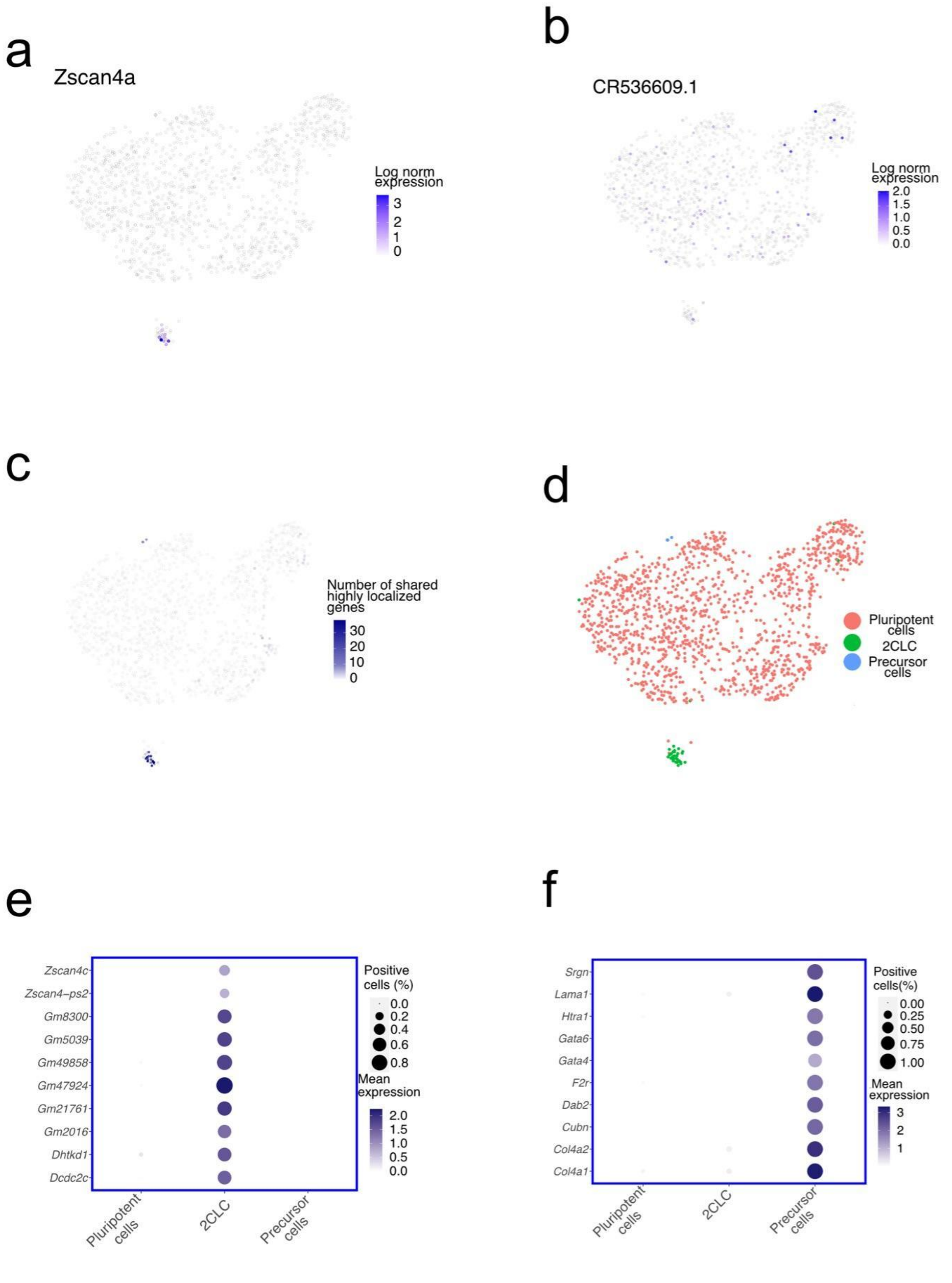


**Supplementary Figure 1**. Application of CIARA to a scRNA-seq data from mouse ESC, taken from [1]. **a**, UMAP representation of the dataset with the expression pattern of a highly localized gene found by CIARA, *Zscan4a.* **b**, Same UMAP representation as in (a) with the expression pattern of a gene not selected by CIARA, *CR536609*.1. **c**, UMAP representation of the mESC dataset indicating the number of highly localized genes expressed by each cell and shared with their neighbours. A greater number of such genes is found in a small group of cells at the bottom, representing a rare population of 2-cell-like cells (2CLC) **d**, UMAP representation of the mESC dataset with the cluster partition found by CIARA**.** Top marker genes of the 2CLC (**e**) and precursor cell (**f**) populations. The size of the dot is given by the fraction of cells with log norm counts above 1 (function NormalizeData from R library Seurat).

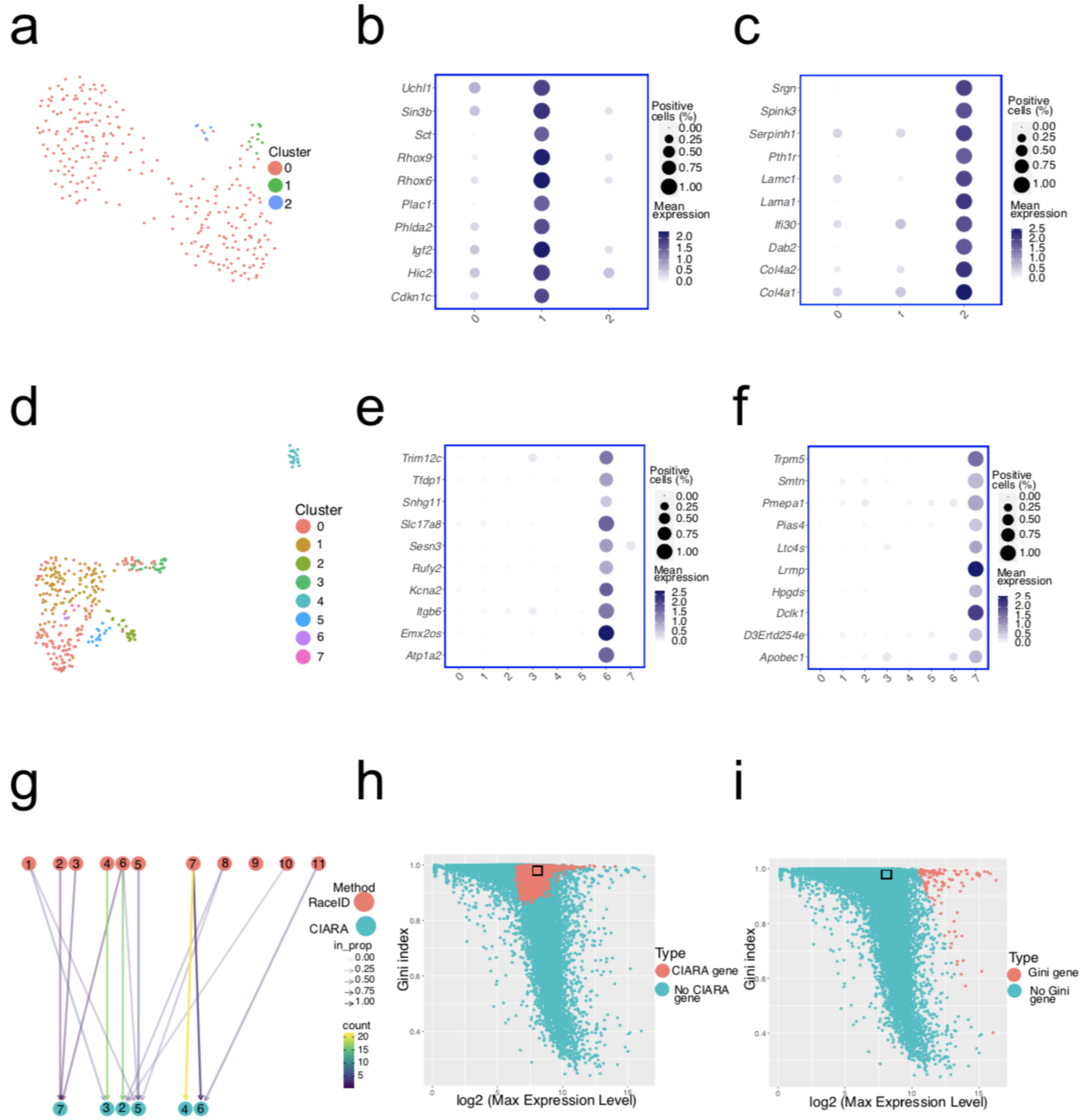


**Supplementary Figure 2**. Analyses of published datasets with CIARA and alternative algorithms. **a**, UMAP representation with the cluster partition found by CIARA in the mouse ESCs scRNA-seq dataset analyzed in the GiniClust2 paper [2]**. b**, **c**, Top marker genes of the two rare populations detected by CIARA in the dataset shown in (a). The size of the dot is given by the fraction of cells with log norm counts above 1 (function NormalizeData from R library Seurat). **d,** UMAP representation of the murine intestinal epithelial cell dataset analyzed in RaceID vignette (https://cran.r-project.org/web/packages/RaceID/vignettes/RaceID.html), with the cluster partition found by CIARA

**e**, **f**, Top marker genes of the two smallest populations (respectively 4 and 3 cells in clusters 6 and 7) detected by CIARA. The size of the dot is given by the fraction of cells with log norm counts above 0.5 (see Method).

In particular, cluster 7 expresses typical markers of Tuft cells [3] **g,** clustree ploty to investigate the relationship between rare clusters found by CIARA (<= 25 cells, top circles) and the original clusters provided by RaceID (bottom circles;  Cluster 2 corresponds to Goblet cells (*Clca3* as marker), cluster 3 to enterocytes (*Apoa1* as marker), cluster 4 to Paneth cells (*Defa24* as marker), cluster 5 to enteroendocrine cells (*Chgb* as marker), cluster 7 to Tuft cells). The clusters found with CIARA correspond to single cell types, while these are split into several clusters with RaceID.

**h**, **i**, Scatterplots of the Gini index as function of the mean expression values for all the genes in the human gastrula dataset [4]. The red circles mark the genes selected by CIARA (panel h) or GiniClust2 (panel i). The black rectangle in the two panels indicate where some of the strongest markers of Primordial Germ Cells are (*NANOS3, NANOG, SOX17* and *DPPA5*).

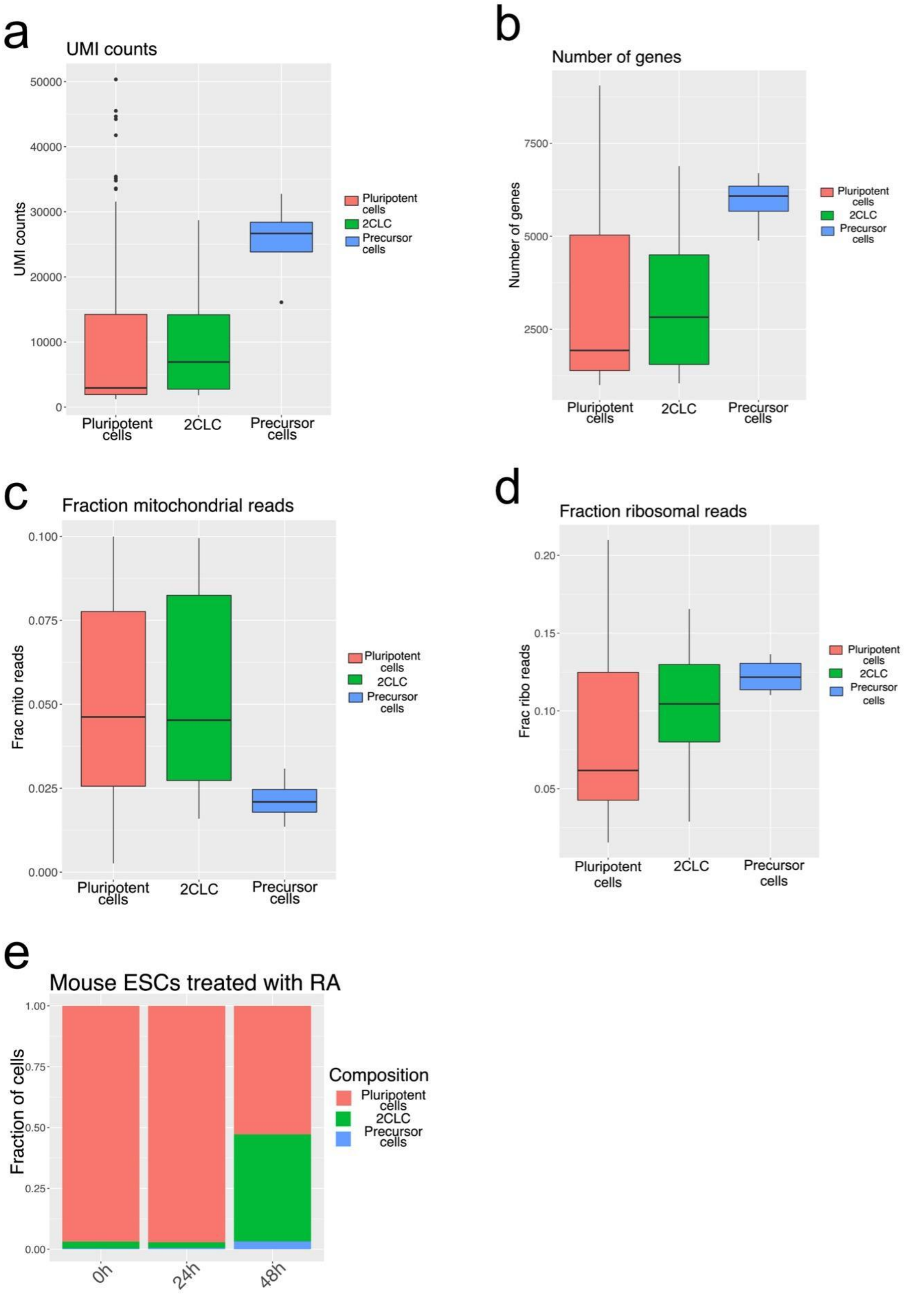


**Supplementary Figure 3.** Analysis of a scRNA-seq dataset from mouse ESCs treated with retinoic acid (RA) for 24h. Boxplot of UMI counts (**a**), number of expressed genes (**b**), fraction of mitochondrial (**c**) and ribosomal reads (**d**). **e**, Cell type composition in mouse ESCs dataset treated with RA for 0, 24 and 48h.

All box plots show the lower quartile (Q1, 25th percentile), the median (Q2, 50th percentile) and the upper quartile (Q3, 75th percentile). Box length refers to interquartile range (IQR, Q3 − Q1). The upper whisker marks the minimum between the maximum value in the dataset and 1.5 times the IQR from Q3 (Q3 + 1.5 × IQR), while the lower whisker marks the maximum between the minimum value in the dataset and the IQR times 1.5 from Q1 (Q1 − 1.5 × IQR). Outliers are shown outside the interval defined by box and whiskers as individual points.

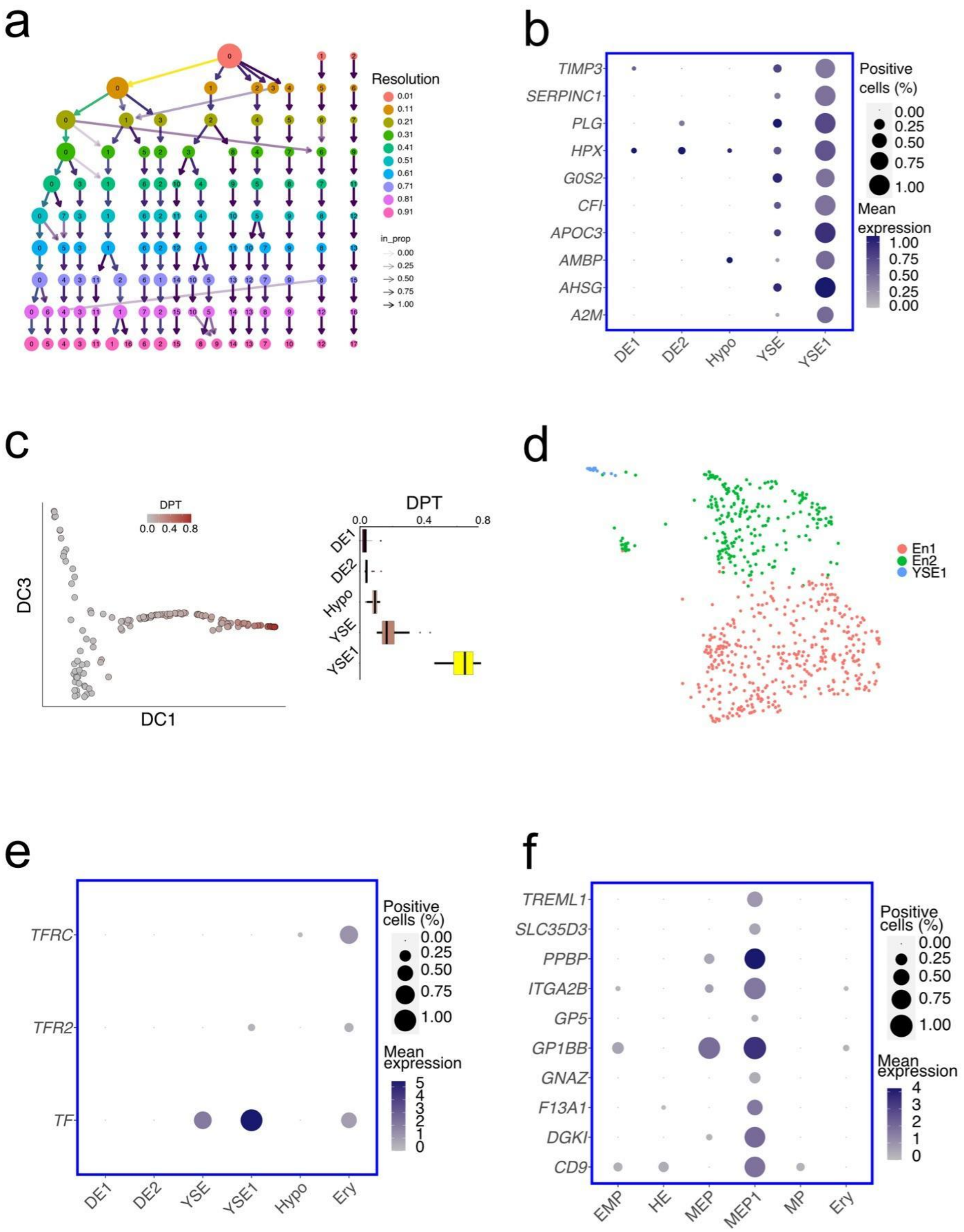


**Supplementary Figure 4.** Analysis of a human gastrula dataset [4]. **a**, clustree plot [5] showing the relationship of the clusters found with a Louvain algorithm for different values of resolution ranging

between 0.01 up to 1. The genes used for clustering are those selected by CIARA. The primordial germ cells cluster (last column on the right) remains unaltered at all values of resolutions.
**b**, Extended list of top markers of the YSE1 cluster. Mean expression levels are normalized by the maximum within each cluster, and the size of the dot is given by the fraction of cells with log norm counts above 1 (function NormalizeData from R library Seurat). **c**, The left panel shows the diffusion components 1 and 3 (DC1 and DC3) of endodermal cells. Cells are colored based on the corresponding value of the diffusion pseudo-time (DPT). The DPT values of cells in each cluster are shown as boxplots in the right panel.
**d**, UMAP representation of the mouse endoderm dataset [6] with the cluster partition found by CIARA. **e**, Balloon plot of transferrin (*TF*) and its two receptors *TFRC* and *TFR2* among endoderm and erythroblasts. The size of the dot is given by the fraction of cells with log norm counts above 1 (see Methods). **f**, Extended list of the top markers of the MEP1. The size of the dot is given by the fraction of cells with log norm counts above 1 (function NormalizeData from R library Seurat).
